# Investigating Cerebral Autoregulation in Traumatic Brain Injury via Simultaneous Measurements of Intracranial Pressure, Arterial Blood Pressure and relative Cerebral Blood Flow

**DOI:** 10.1101/2023.02.10.528028

**Authors:** Christopher Uff

## Abstract

Traumatic Brain Injury (TBI) can alter the brain’s ability to maintain an adequate supply of oxygen and metabolites to brain tissue by disrupting the autoregulatory mechanisms that maintain constant cerebral blood flow. Impaired cerebral autoregulation can result in brain hypoxia leading to morbidity and mortality so maintenance of cerebral blood flow after injury is of paramount importance. Currently, this is managed using limited information and various assumptions, hence there is significant interest in developing better correlates for establishing whether cerebral autoregulation is impaired or intact. In this study we simultaneously measure cerebral blood flow (CBF) using a non-invasive optical approach, intracranial pressure (ICP, measured invasively) and arterial blood pressure (ABP) with the aim of investigating the relationships between these signals over multiple timescales, and ultimately assessing how these measurements may best be combined and interpreted to aid the treatment of TBI.

## Introduction

Cerebral Autoregulation (CA) is the complex process whereby the body maintains constant blood flow and therefore constant oxygenation and nutrition provision to the brain. By balancing blood and cerebrospinal fluid (CSF) pressures, CA dynamically stabilises cerebral perfusion pressure (CPP) and hence cerebral blood flow^1^.

CA encompasses a broad set of changes to vascular physiology that involves several tissues and cell types^2^. The CA *myogenic mechanism* maintains constant blood flow in the face of changing blood pressure. In response to an increase in ABP, autoregulatory mechanisms sense the stretch in cerebral arteries and arterioles and translate this to a constriction of surrounding arterial smooth muscle, resulting in increased cerebrovascular resistance to offset increased ABP (and lower stretch results in relaxation to offset decreased ABP). In contrast, using the *metabolic mechanism*, CA senses local changes in blood CO_2_, and acts on arterial smooth muscle via nitric oxide pathways causing dilation or constriction, thus changing blood perfusion according to the local demands of brain metabolism. Additional mechanisms involve neurons or endothelial cells that through a variety of signalling mechanisms can act on smooth muscle tone. The objectives of these mechanisms are similar; to maintain constant cerebral blood flow and thus constant delivery of oxygen and metabolites to the brain in the face of fluctuating arterial blood pressure and cerebral metabolic demand ^1,2^.

Failure of CA is frequently seen in severe brain injury and can result in reduced oxygen and nutrient delivery leading to significant morbidity and mortality. By different measures, CA is impaired in 49–87% of TBI patients.^3,4^ However it is difficult to ascertain if CA impairment underlies the physiological trends observed clinically. For instance, cerebral hypoperfusion may result from mismatched metabolic demand resulting in loss of consciousness. Ischaemic tissue damage may be compounded by impaired CA resulting in infarction of critically ischaemic tissue. Monitoring cerebral perfusion is thus a significant part of modern neurointensive care.

Clinical decision making related to CA status is currently informed by mismatches in ICP responses to blood pressure variations: if ICP increases with an increase in cerebral blood pressure, it is assumed that CA is not intact (intact CA maintains constant cerebral blood flow and hence ICP as Bp increases). It remains unresolved whether additional information is available from the relationships between cerebral blood flow, arterial blood pressure, and intracranial pressure, and in what clinical conditions this information is most useful.

Patients with severe TBI are likely to have disrupted CA, but there is at present no widely used method for measuring cerebral perfusion that is non-invasive, accurate, and able to be used continually. The best available method to continuously monitor CBF and its relationship with systemic blood pressure is trans-cranial doppler ultrasound^2^ which lacks consistency, is operator dependent and only measures blood flow in major arteries (for example middle cerebral artery) and therefore lacks the ability to measure microvascular blood flow in brain tissue.

Here, we employ a non-invasive measurement of CBF using Near-Infra-Red Diffuse Correlation Spectroscopy (NIR-DCS) to monitor brain perfusion and CA. We present case studies that are the first of their kind in the UK to combine these signals in the context of traumatic brain injury. We establish that in these subjects, NIR-DCS signal quality is sufficient to extract indices related to the autoregulatory state of the brain, and to permit further investigation into the utility of using non-invasive blood flow measurements in the clinic.

## Methods

### Ethics

Subjects were recruited in the critical care unit of the Royal London Hospital, London in accordance with a protocol approved by a UK Health Research Authority (IRAS 317751). Ethical approval was obtained from the London - South East Research Ethics Committee (REC reference: 22/LO/0825) on 15/11/22. This protocol allowed for the recruitment of patients undergoing invasive ICP monitoring as a routine part of their care. The dominant group of patients in this category at The Royal London hospital are Traumatic Brain Injury patients. All study procedures were carried out in accordance with the ethical principles in the Research Governance Framework for Health and Social Care, Second Edition, 2005. Conscious patients gave written and informed consent prior to any data collection. In the case of unconscious patients, assent to participation was obtained from next of kin acting as a personal consultee or from an independent healthcare professional acting as a professional consultee according to the UK Mental Capacity Act and the Declaration of Helsinki^5^.

### Measurement

Measurements of cerebral blood flow were obtained using a custom near-infrared optical research instrument (NIR-DCS) provided by CoMind Technologies, London. The instrument contains a laser source and a set of optical detectors that form a single non-invasive measurement on the forehead. The laser source was regulated to keep its optical output within skin-safety limits defined by established international safety standards (IEC 60825-1:2014). All electronics were housed within a medical-grade cart that met all relevant electrical safety standards (IEC-60601-1). Optical fibres were used to carry light from the laser source to a head probe that was placed on the participant’s forehead. Light that scatters through the subject’s brain was collected by optical fibres integrated into the head-probe, which carried that light to optical detectors. Stable positioning on the subject’s forehead was ensured using medical-grade tape (3M Medical Tape 1509).

Subjects were included in the study if they were undergoing invasive ICP monitoring alongside arterial-line measurements of ABP as standard medical care. Additional exclusion criteria included subjects with radiological evidence of a severely compromised skull (including via decompressive hemicraniectomy), and subjects with machine-regulated cardiac output (e.g. Cardiopulmonary Bypass or Arterio-Venous ECMO).

Arterial Blood Pressure was measured invasively using an arterial catheter interfaced to a transducer for the duration of routine clinical monitoring. Intracranial pressure was measured invasively using an intraparenchymal pressure transducer (Raumedic Neurovent-P). Both signals were collected and monitored continuously through a clinical patient monitor (Mindray BeneVision N1). For the purposes of research data collection, ICP and CBF were duplicated as analog voltages sent from the Mindray and read into a custom digitiser device, sampled at 100 Hz and synchronized with blood flow signals from the NIRS-DCS device.

### Analysis

Cerebral Blood Flow was estimated using NIR-DCS data by fitting the unnormalized temporal electric field autocorrelation decay of detected light to a model of the correlation diffusion equation under the assumption that scatterer dynamics follow Brownian motion^6,7^. After incorporating constant values for tissue reduced scattering and absorption (□a: 0.1 cm^-1^, □s’: 9.0 cm^-1^), this model outputs a particle diffusion coefficient (*D*_*B*_ in mm^2^/s), which is proportional to CBF^8^.

Analog signals representing ICP, ABP, were digitised at 100 Hz whilst CBF was digitised at 20 Hz. Raw patient data was saved to a hard-drive, transferred to a secure workstation located within the Royal London Hospital, stored according to best-practice clinical data security and GDPR regulations. These data were analysed on-site using custom software written in Python.

## Results

Measurements were obtained from subjects admitted to the hospital for treatment following Traumatic Brain Injury (TBI). Sixteen patients have been studied to date. A summary and examples of pertinent patient data is provided in Table 1. In all subjects, ICP and ABP were monitored continuously using invasive probes as part of routine clinical monitoring. In addition to routine monitoring, the same ICP and ABP signals were captured alongside CBF using the NIR-DCS research instrument in a continuous fashion over a number of hours in each patient. Recording was only interrupted if the patient had to be moved (for example, for a CT scan), or for other clinical reasons.

**Table 1:**
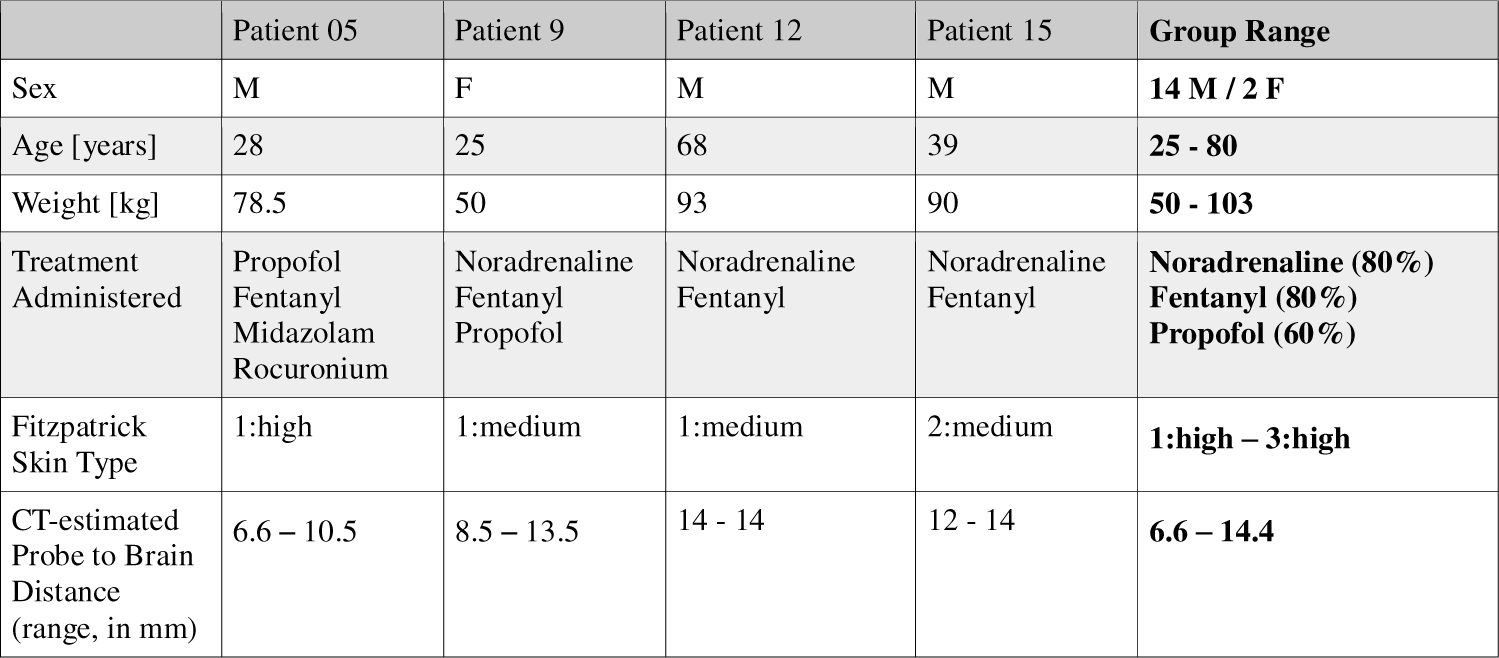
Summary of pertinent patient data. Personal and medical history information was collected on all patients. This table shoes a sub-sample of this data and group summary ranges. More complete patient data are available on reasonable request.

|  | Patient 05 | Patient 9 | Patient 12 | Patient 15 | Group Range |
| --- | --- | --- | --- | --- | --- |
| Sex | M | F | M | M | 14 M / 2 F |
| Age [years] | 28 | 25 | 68 | 39 | 25 - 80 |
| Weight [kg] | 78.5 | 50 | 93 | 90 | 50 - 103 |
| Treatment Administered | Propofol<br>Fentanyl<br>Midazolam<br>Rocuronium | Noradrenaline<br>Fentanyl<br>Propofol | Noradrenaline<br>Fentanyl | Noradrenaline<br>Fentanyl | Noradrenaline (80%)<br>Fentanyl (80%)<br>Propofol (60%) |
| Fitzpatrick Skin Type | 1:high | 1:medium | 1:medium | 2:medium | 1:high – 3:high |
| CT-estimated Probe to Brain Distance (range, in mm) | 6.6 – 10.5 | 8.5 – 13.5 | 14 - 14 | 12 - 14 | 6.6 – 14.4 |

The data quality of ICP and ABP data was consistently high and resulted in well-resolved pulsatile waveforms resting on top of slower physiological fluctuations. The CBF signals (as measured via NIRS-DCS) were noisier, as is expected for non-invasive optical measurements, but nonetheless showed clear cardiac pulsatility on top of slower fluctuations in all patients. Examples are provided in **Figure 1A and Figure 1B**. Full time-series plots are provided for each subject in Supplementary Data. The data presented in this paper is available on reasonable request.

**Figure 1:**
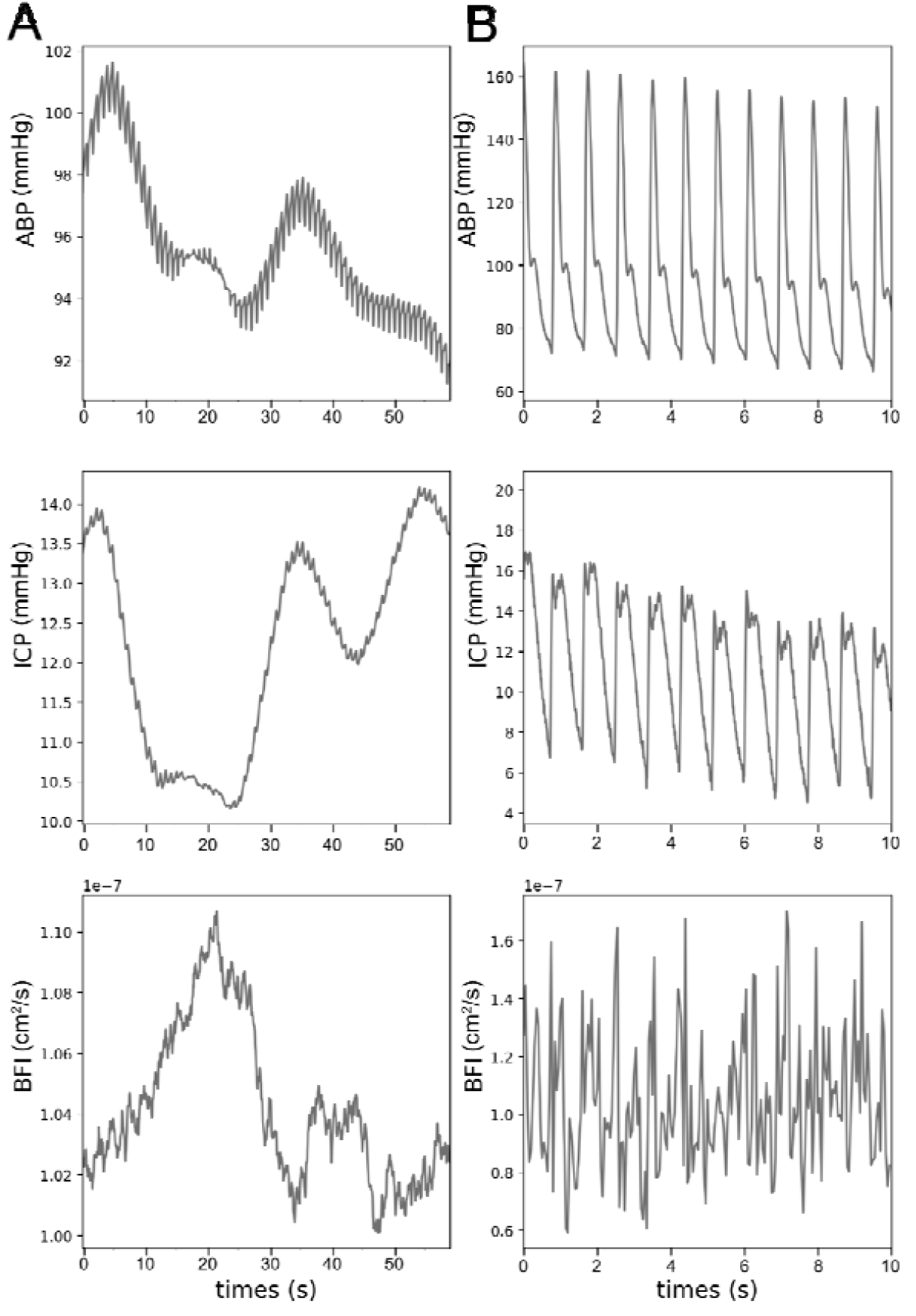
Example time-series of data collected in this study. Arterial Blood Pressure (ABP), Intracranial Pressure (ICP) and Cerebral Blood Flow (CBF) were simultaneously measured for each subject. A) Comparison of CBF, ABP and ICP on the time-scale of minutes. Raw traces are filtered using a moving average applied over segments of 10 seconds. B) Comparison of raw pulsatile waveforms from CBF, ABP and ICP.

Intact cerebral autoregulation buffers CBF from fluctuations in ABP. To test if this was evident in our data, we correlated ABP and CBF, which we have termed the Flow Reactivity Index (FRx) to determine if there was any relationship between them. For example, in Subject 1 we observed a positive correlation (Pearson’s R^2^=0.63) between CBF and ABP as is evident in **Figure 2A**. This compares to the weaker correlation observed between CBF and ICP (Pearson’s R^2^=0.36, **Figure 2B**).

**Figure 2:**
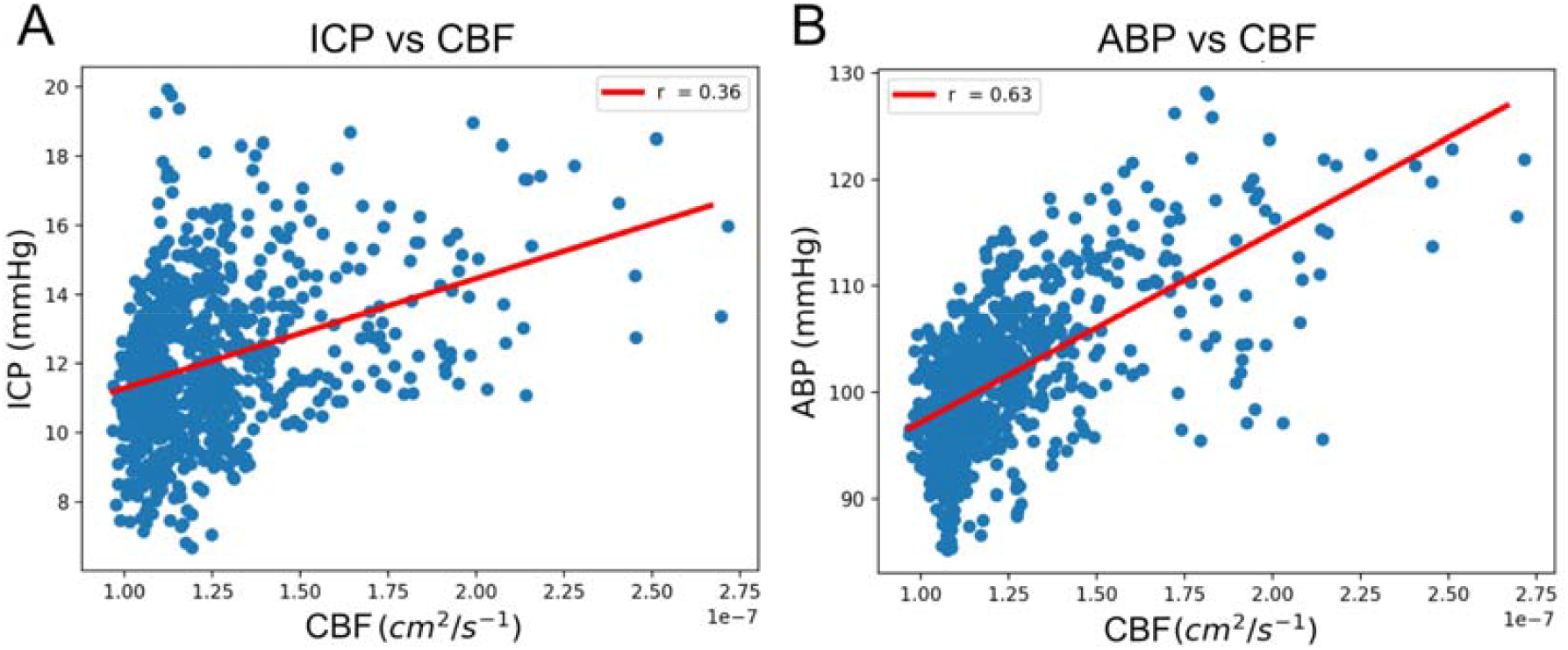
Correlation-based measurements of CA. A) Correlation between ICP and ABP. B) Correlation between CBF and ABP.

## Discussion

We have presented a case series that demonstrates that it is possible to investigate cerebral blood flow using non-invasive optical measures alongside invasive clinical measurements of ABP and ICP. NIR-DCS data were of sufficient quality to identify expected pulse waveform features of CBF representing the characteristic cardiac pressure waves, and measurements over multiple timescales showed correlated fluctuations between the three signals. Together, these data indicate that it is now technically possible to simultaneously and continuously measure ABP, BFI and ICP at the bedside in TBI patients, potentially unlocking significant clinical utility through the analysis and interpretation of these critical clinical variables.

Whether Cerebral Autoregulation mechanisms are intact or not can be indicated by the cerebrovascular pressure reactivity (the ability of vascular smooth muscle to respond to changes in cerebral perfusion pressure). This has been previously measured by relating ABP to ICP to produce the Pressure Reactivity Index (PRx), or by relating ABP to transcranial doppler to measure the mean-flow index (Mx)^9^, to laser speckle doppler to produce a laser doppler index (LDx)^10^, or to cerebral oximetry to produce a cerebral oximetry index (COx)^11^. In this study, we directly compare measurements of CA using pressure reactivity and flow reactivity, and found in preliminary analysis that significant correlations between BFI exist with both ABP and ICP.

By increasing this case series and collecting many consecutive hours of data per subject, it may be possible to estimate the optimal CPP by establishing a ‘U-shaped’ relationship between PRx and CPP^12^. Finding a similar minimisation in the relationship between FRx and CPP would potentially open new avenues of investigation for using CBF measurements to guide clinical decision making.

## References

1. Panerai RB. Cerebral Autoregulation: From Models to Clinical Applications. Cardiovasc Eng. 2008;8(1):42–59. doi:10.1007/s10558-007-9044-6

2. Armstead WM. Cerebral Blood Flow Autoregulation and Dysautoregulation. Anesthesiol Clin. 2016;34(3):465–477. doi:10.1016/j.anclin.2016.04.002

3. Hlatky R, Valadka AB, Robertson CS. Intracranial Pressure Response to Induced Hypertension: Role of Dynamic Pressure Autoregulation. Neurosurgery. 2005;57(5):917–923. doi:10.1227/01.NEU.0000180025.43747.fc

4. Bouma GJ, Marmarou A. Blood pressure and intracranial pressure-volume dynamics in severe head injury: relationship with cerebral blood flow.

5. Puri KS, Suresh KR, Gogtay NJ, Thatte UM. Declaration of Helsinki, 2008: implications for stakeholders in research. J Postgrad Med. 2009;55(2):131–134. doi:10.4103/0022-3859.52846

6. Boas DA, Campbell LE, Yodh AG. Scattering and Imaging with Diffusing Temporal Field Correlations. Phys Rev Lett. 1995;75(9):1855–1858. doi:10.1103/PhysRevLett.75.1855

7. Verdecchia K, Diop M, Morrison LB, Lee TY, St. Lawrence K. Assessment of the best flow model to characterize diffuse correlation spectroscopy data acquired directly on the brain. Biomed Opt Express. 2015;6(11):4288. doi:10.1364/BOE.6.004288

8. Durduran T, Choe R, Baker WB, Yodh AG. Diffuse optics for tissue monitoring and tomography. Rep Prog Phys. 2010;73(7):076701. doi:10.1088/0034-4885/73/7/076701

9. Budohoski KP, Czosnyka M, de Riva N, et al. The Relationship Between Cerebral Blood Flow Autoregulation and Cerebrovascular Pressure Reactivity After Traumatic Brain Injury. Neurosurgery. 2012;71(3):652–661. doi:10.1227/NEU.0b013e318260feb1

10. Lam JMK, Hsiang JNK, Poon WS. Monitoring of autoregulation using laser Doppler flowmetry in patients with head injury. J Neurosurg. 1997;86(3):438–445. doi:10.3171/jns.1997.86.3.0438

11. Brady KM, Lee JK, Kibler KK, Easley RB, Koehler RC, Shaffner DH. Continuous Measurement of Autoregulation by Spontaneous Fluctuations in Cerebral Perfusion Pressure: Comparison of 3 Methods. Stroke. 2008;39(9):2531–2537. doi:10.1161/STROKEAHA.108.514877

12. Steiner LA, Czosnyka M, Piechnik SK, et al. Continuous monitoring of cerebrovascular pressure reactivity allows determination of optimal cerebral perfusion pressure in patients with traumatic brain injury: Crit Care Med. 2002;30(4):733–738. doi:10.1097/00003246-200204000-00002

