## Supplementary data S3 - S8 for "Investigating Cerebral Autoregulation in Traumatic Brain Injury via Simultaneous Measurements of Intracranial Pressure, Arterial Blood Pressure and relative Cerebral Blood Flow"

**Professor Christopher Uff***

Professor of Neurosurgery and Head of Neurotrauma, Royal London Hospital, London E1 1FR, UK

* Author to whom correspondence should be addressed.

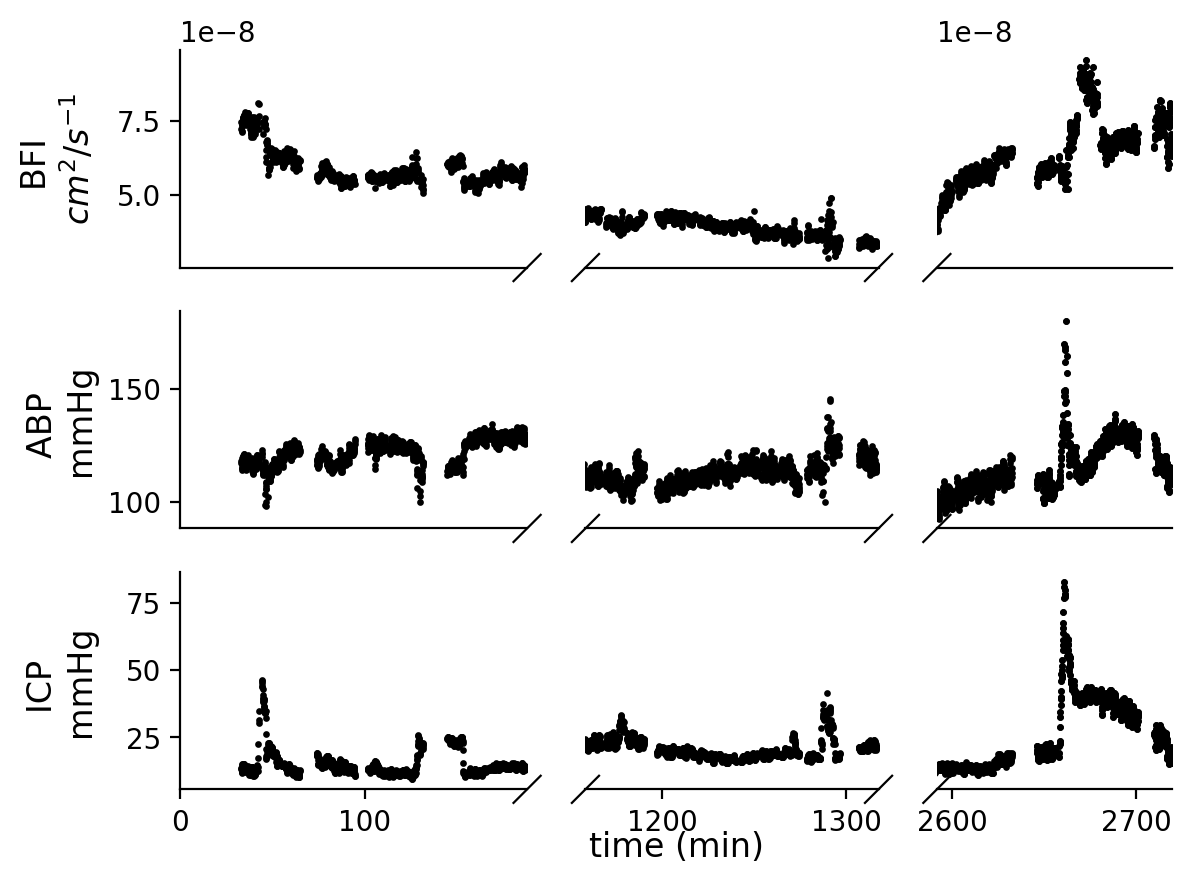

Figure S2. Patient 3.

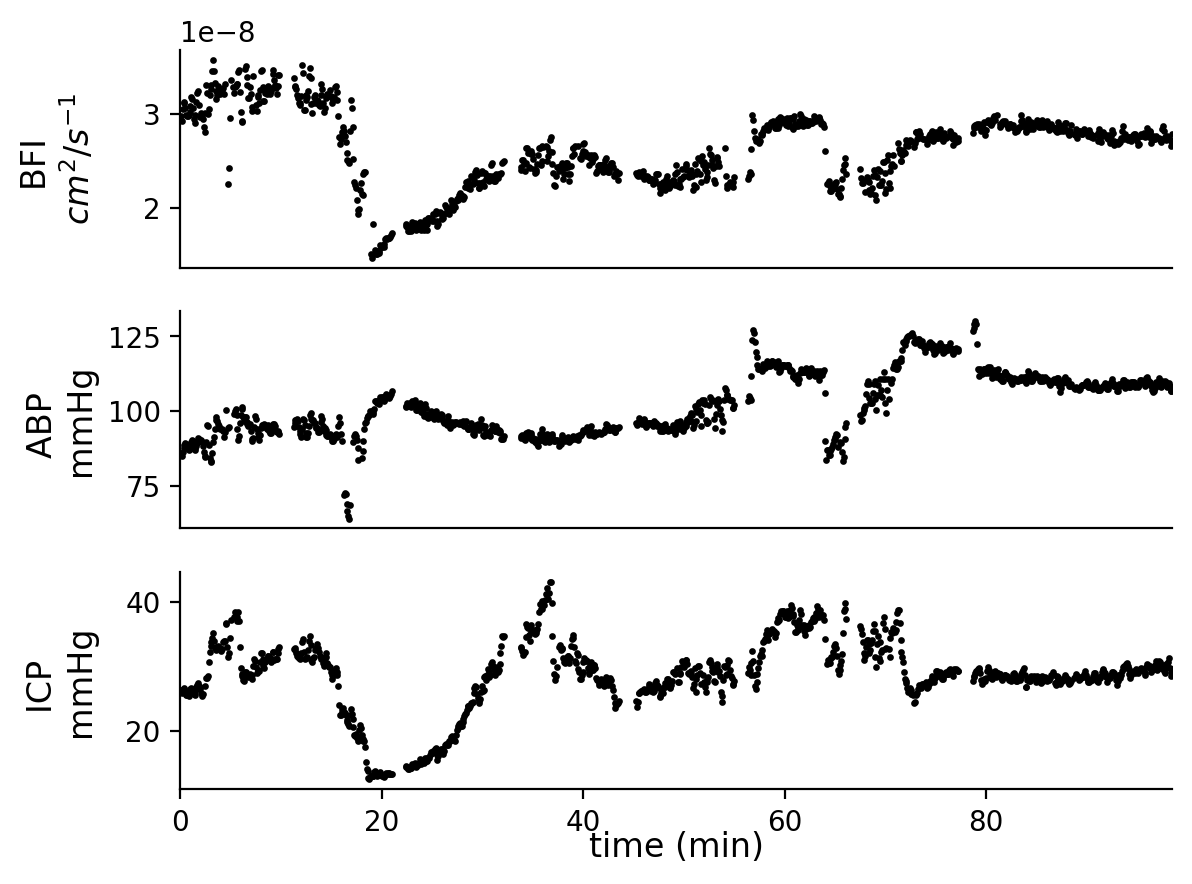

Figure S3. Patient 4.

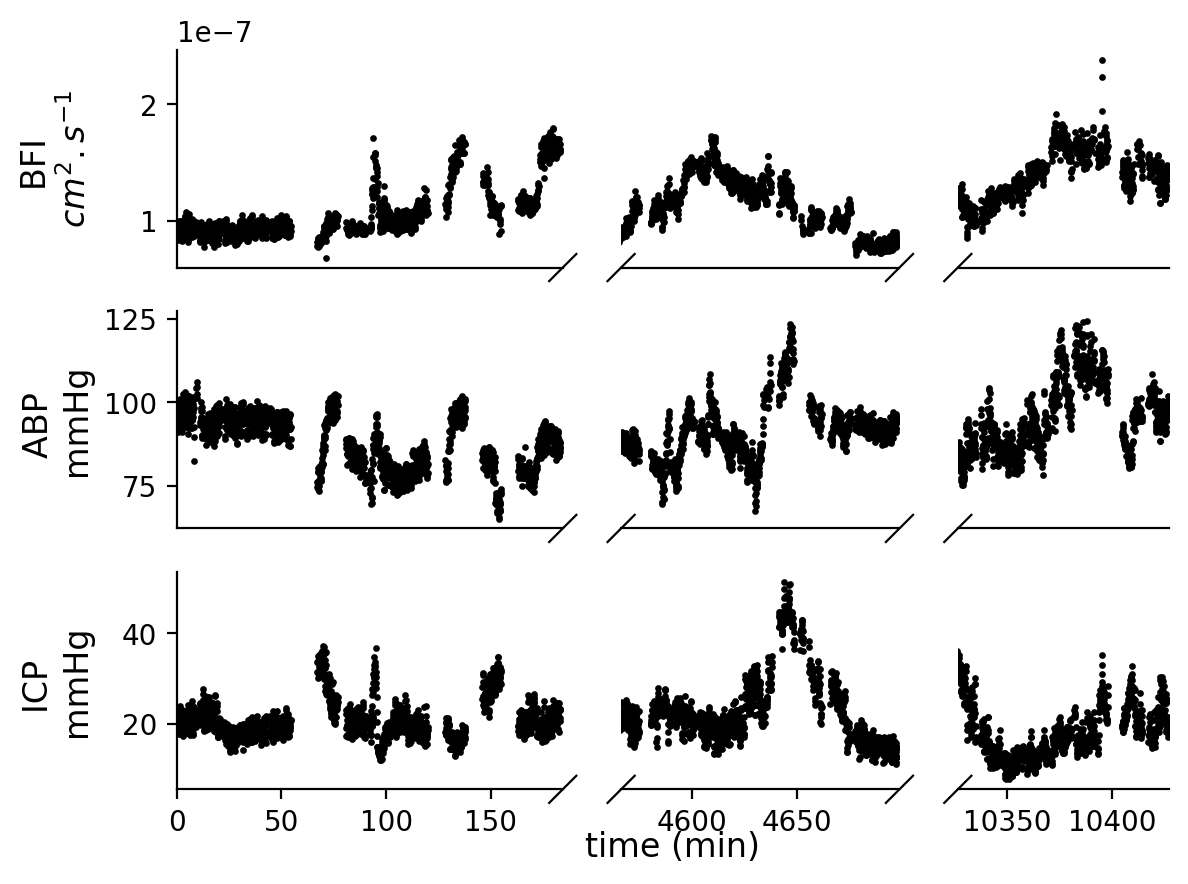

Figure S4. Patient 5.

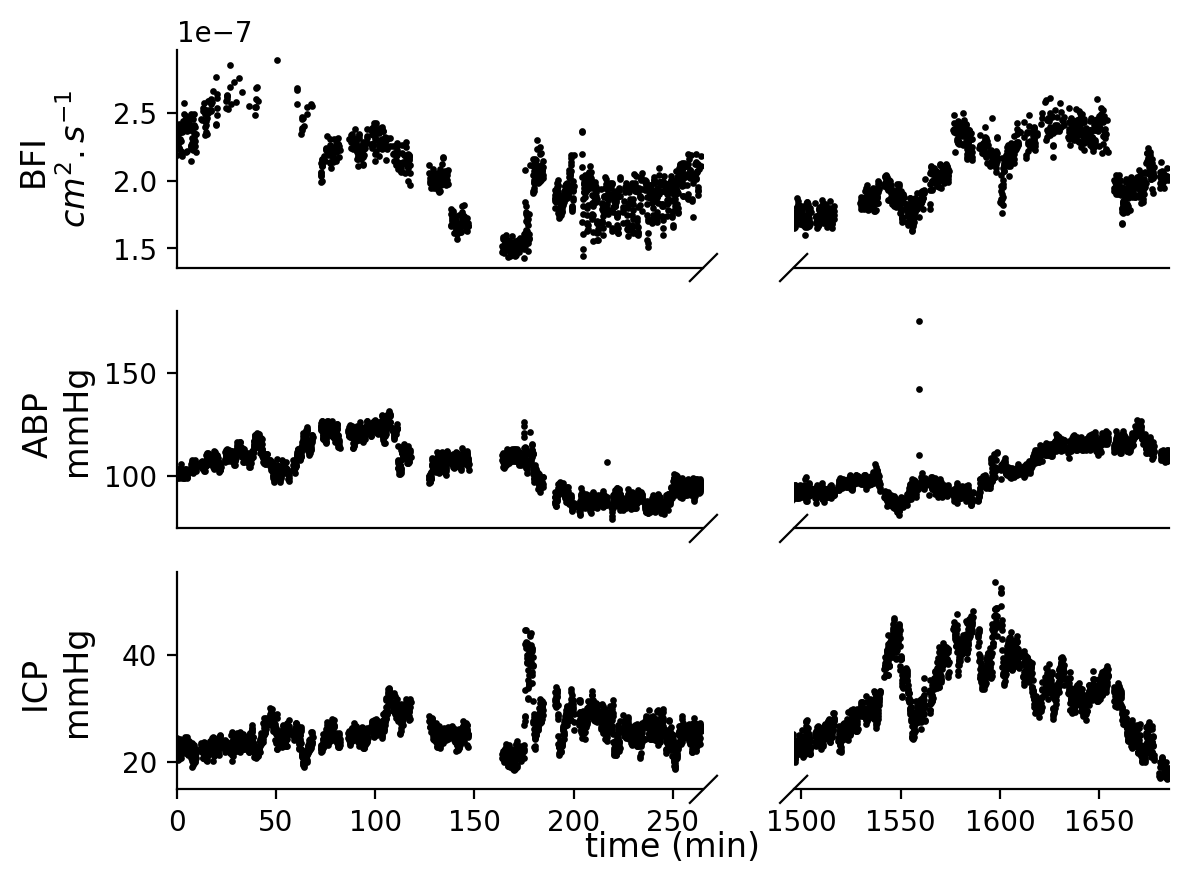

Figure S5. Patient 6.

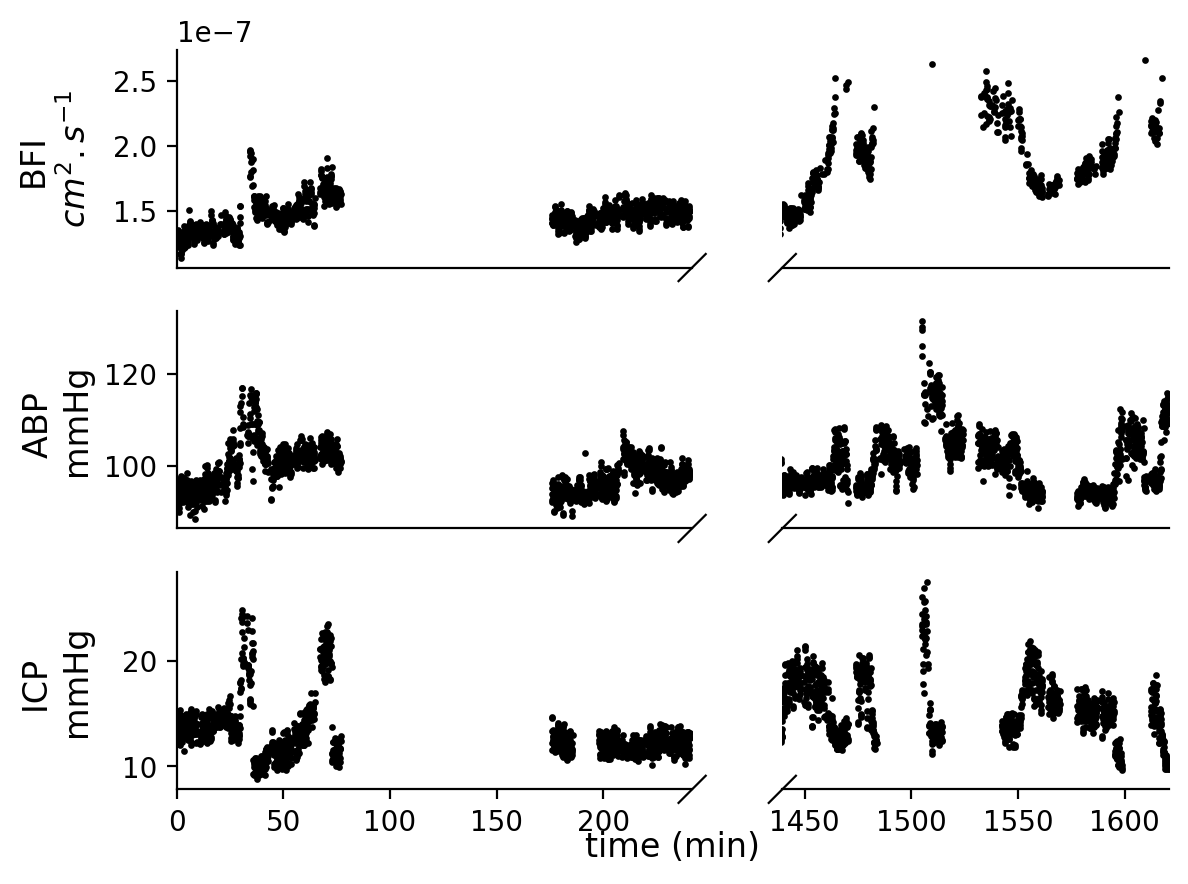

Figure S6. Patient 7.

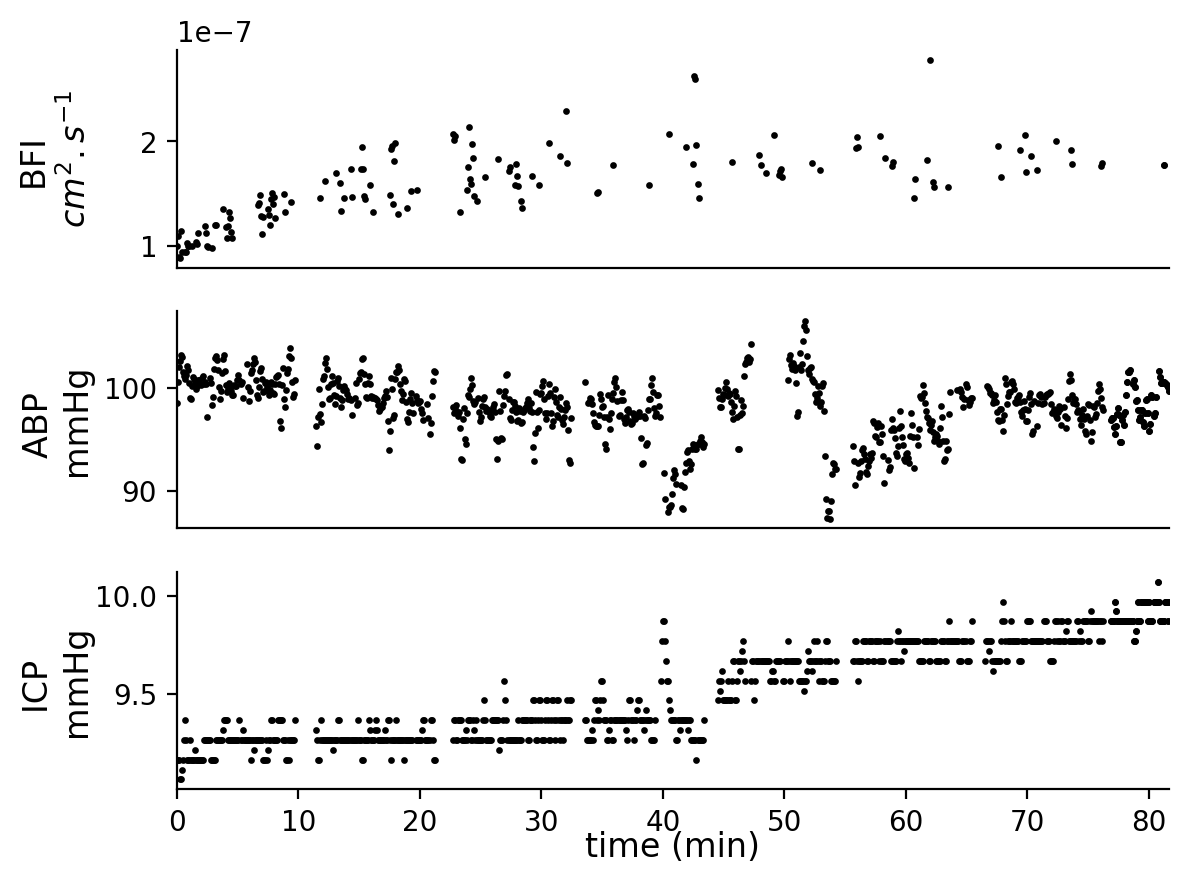

Figure S7. Patient 9.

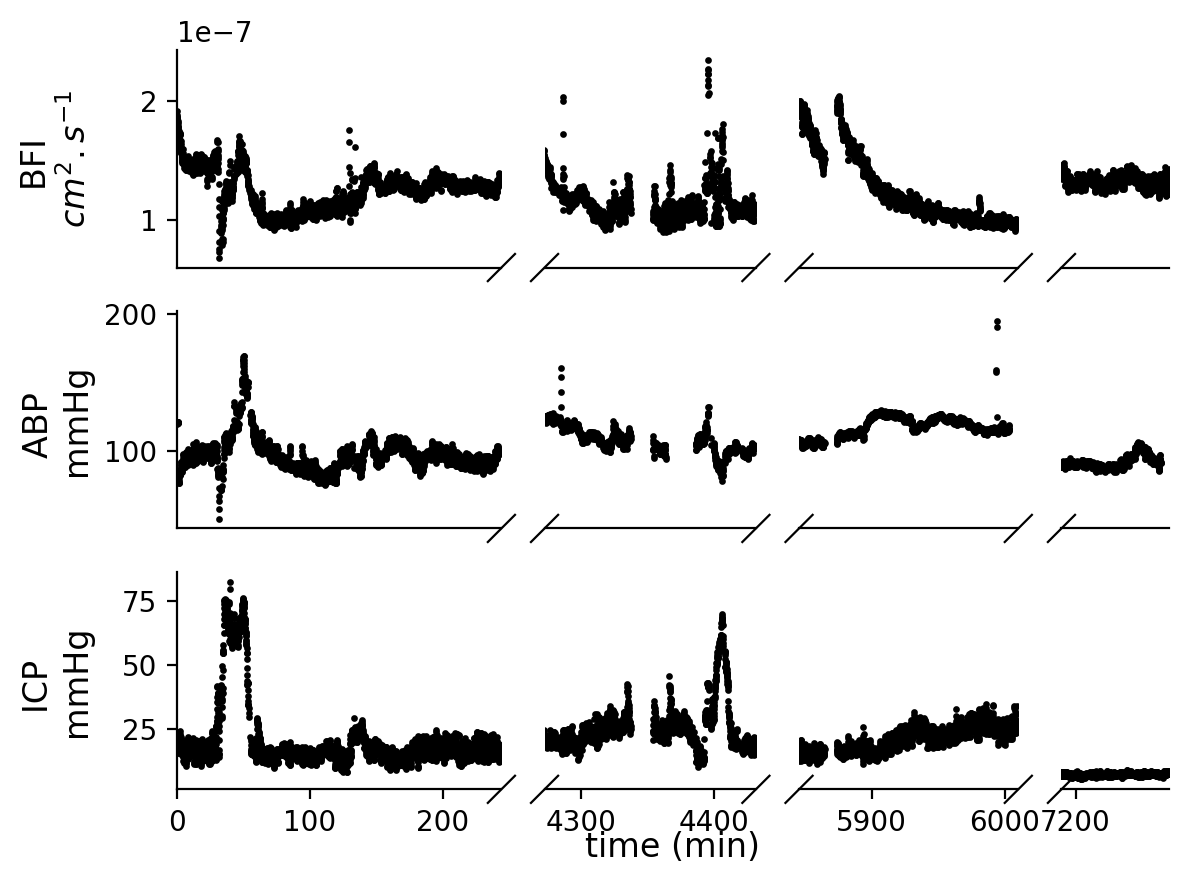

Figure S8. Patient 10.

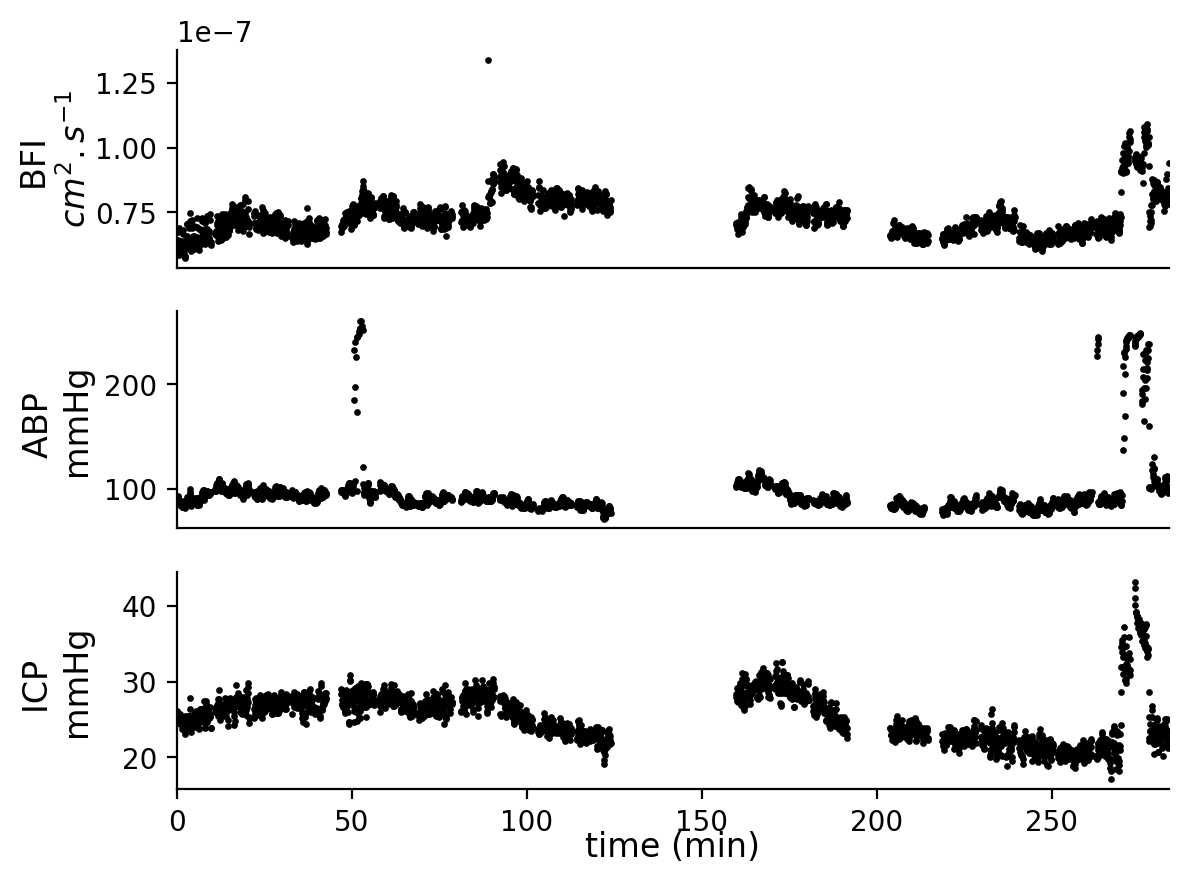

Figure S9. Patient 11.

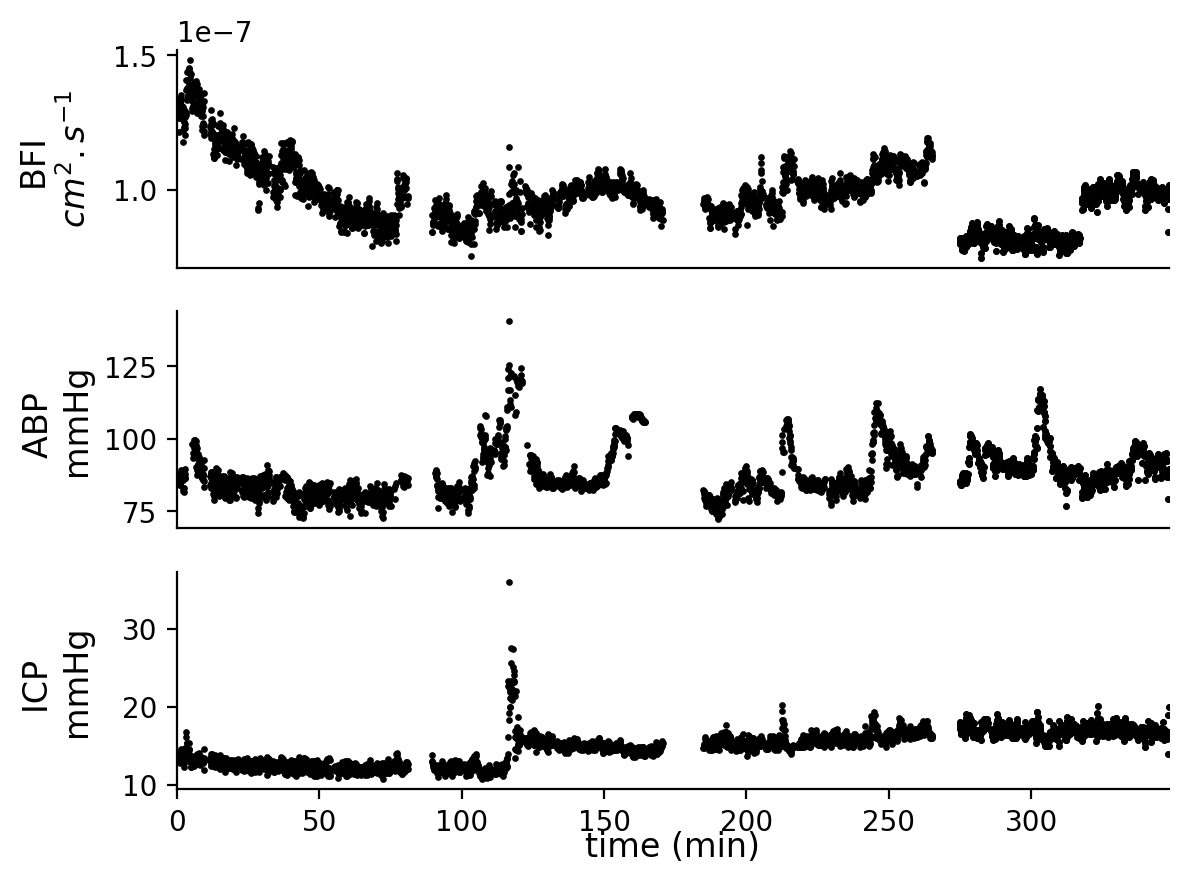

Figure S10. Patient 12.

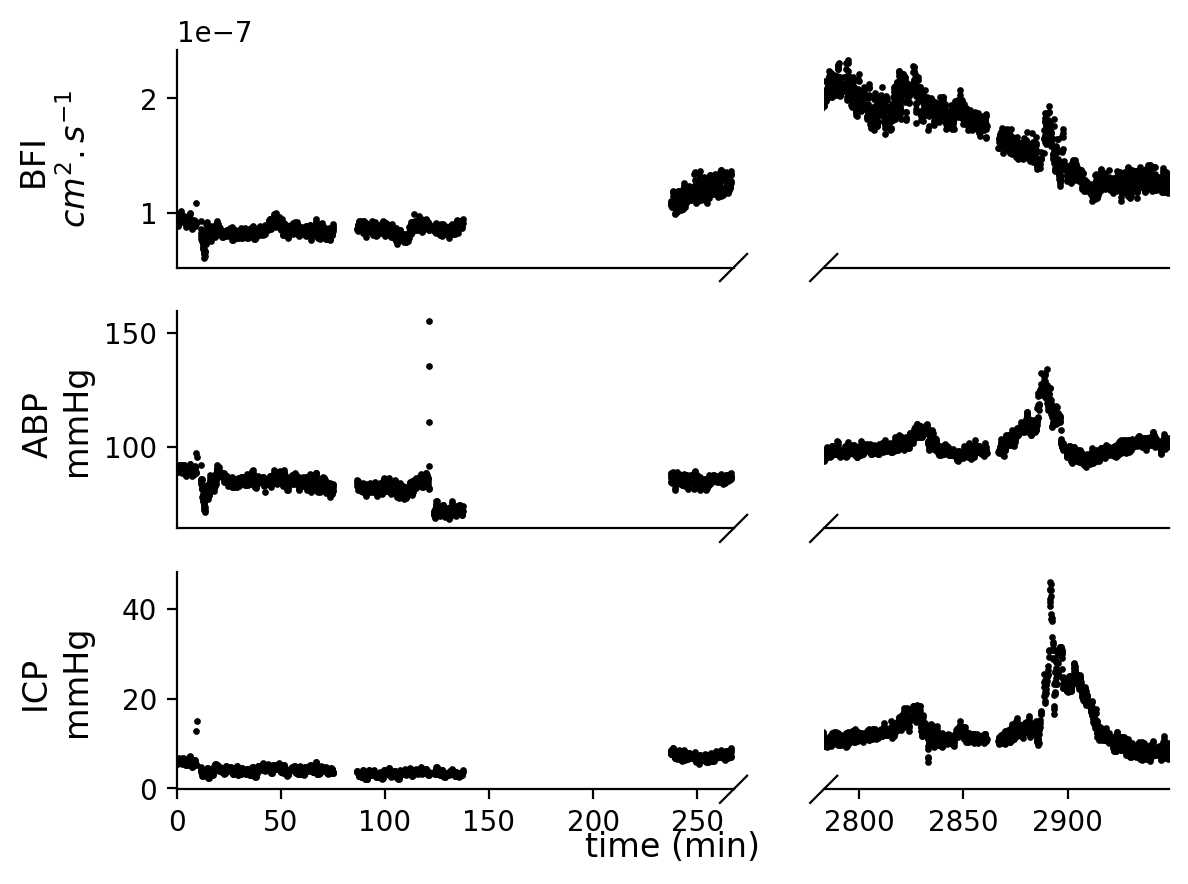

Figure S11. Patient 13.

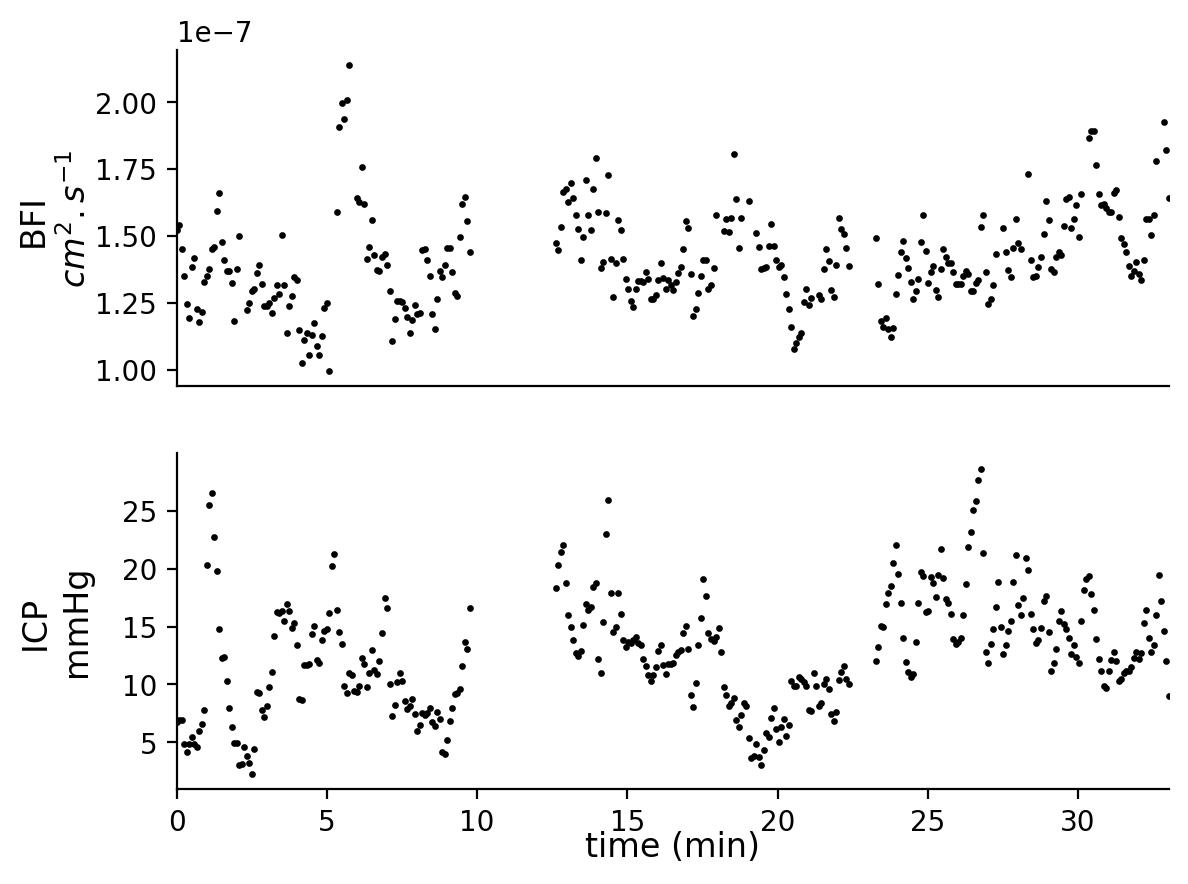

Figure S12. Patient 14.

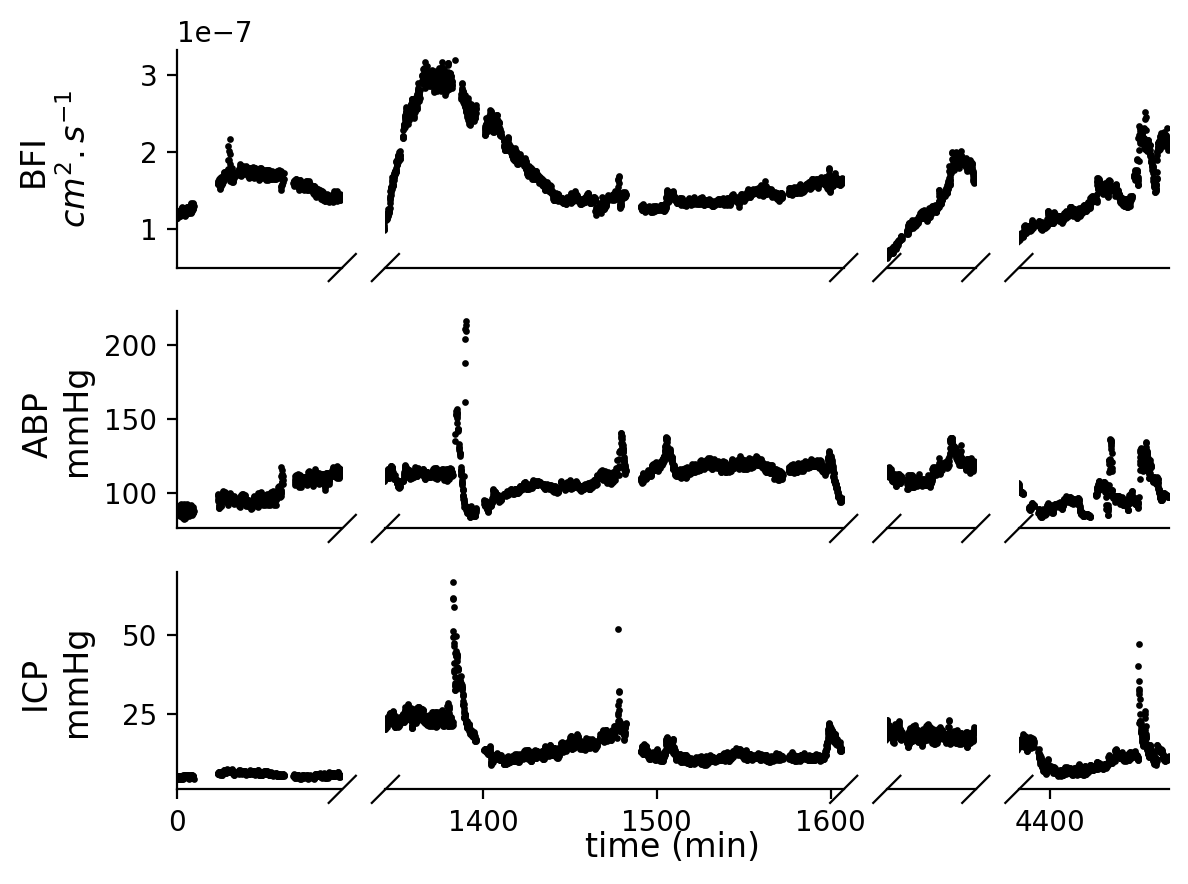

Figure S13. Patient 15.

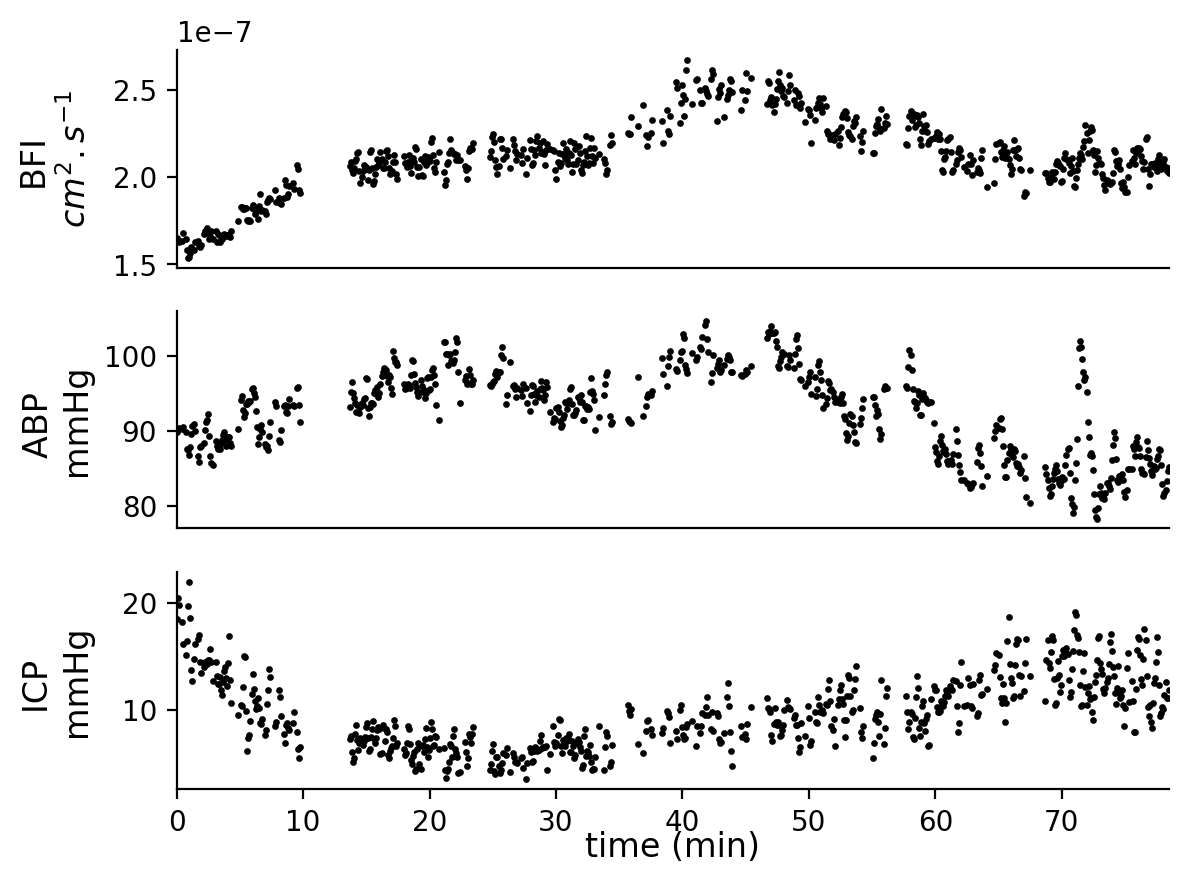

Figure S14. Patient 16.
